# CRISPR-based DNA and RNA detection with liquid-liquid phase separation

**DOI:** 10.1101/471482

**Authors:** Willem Kasper Spoelstra, Jeroen M. Jacques, Franklin L. Nobrega, Anna C. Haagsma, Marileen Dogterom, Timon Idema, Stan J. J. Brouns, Louis Reese

## Abstract

The ability to detect specific nucleic acid sequences allows for a wide range of applications including identification of pathogens, clinical diagnostics, and genotyping. CRISPR-Cas proteins Cas12a and Cas13a are RNA-guided endonucleases that bind and cleave specific DNA and RNA sequences, respectively. After recognition of a target sequence both enzymes activate a unique, indiscriminate nucleic acid cleavage activity, which has been exploited for detection of sequence specific nucleotides using labelled reporter molecules. We here present a label-free detection approach that uses a readout based on solution turbidity caused by liquid-liquid phase separation (LLPS). Turbidity arises from coacervates of positively charged polyelectrolytes with long poly(dT) or poly(U) oligonucleotides. In the presence of a target sequence, long oligonucleotides are progressively shortened, changing the solution from turbid to transparent. We explain how oligonucleotide cleavage resolves LLPS by using a mathematical model which we validate with poly(dT) phase separation experiments. The deployment of LLPS complements CRISPR-based molecular diagnostic applications and facilitates easy and low-cost nucleotide sequence detection.

## Introduction

CRISPR-Cas technology has revolutionized molecular biology by making genome editing (*1*) and nucleic acid sequence detection efficient and accessible (*2*–*6*). CRISPR-based detection methods rely on CRISPR RNA-guided nucleases such as Cas12a (*7*) (type V), Cas14a (*8*), and Cas13a (*9*) (type VI). These proteins display non-specific cleavage activity of ssDNA (Cas12a (*5, 10*) and Cas14a (*8*)) or RNA (Cas13a (*2, 11, 12*)) upon recognition of their target sequences. The mechanisms employed by these enzymes rely on the formation of a Cas protein complex with CRISPR RNA (crRNA), which then targets DNA or RNA sequences complementary to the crRNA (*12*–*14*). Importantly, the enzymes display non-specific (collateral) cleavage activity as long as a target is bound to the crRNA-protein complex (**Fig. 1**). This collateral activity is the mechanism that facilitates nucleotide detection assays in solution or on paper-based tests (*2*–*5*). Previously, Cas12a and Cas13a have been employed to discriminate between two variants of the human papillomavirus (*5*) and between Dengue and Zika viruses (*3*), respectively. In these studies, the detection of Cas12a and Cas13a activity relied on quenched fluorophores linked by DNA or RNA molecules. The proteins’ collateral endonuclease activity, which is activated after target binding, causes a fluorescent signal by cleaving the DNA or RNA linkage between a fluorophore-quencher pair.

**Figure 1:**
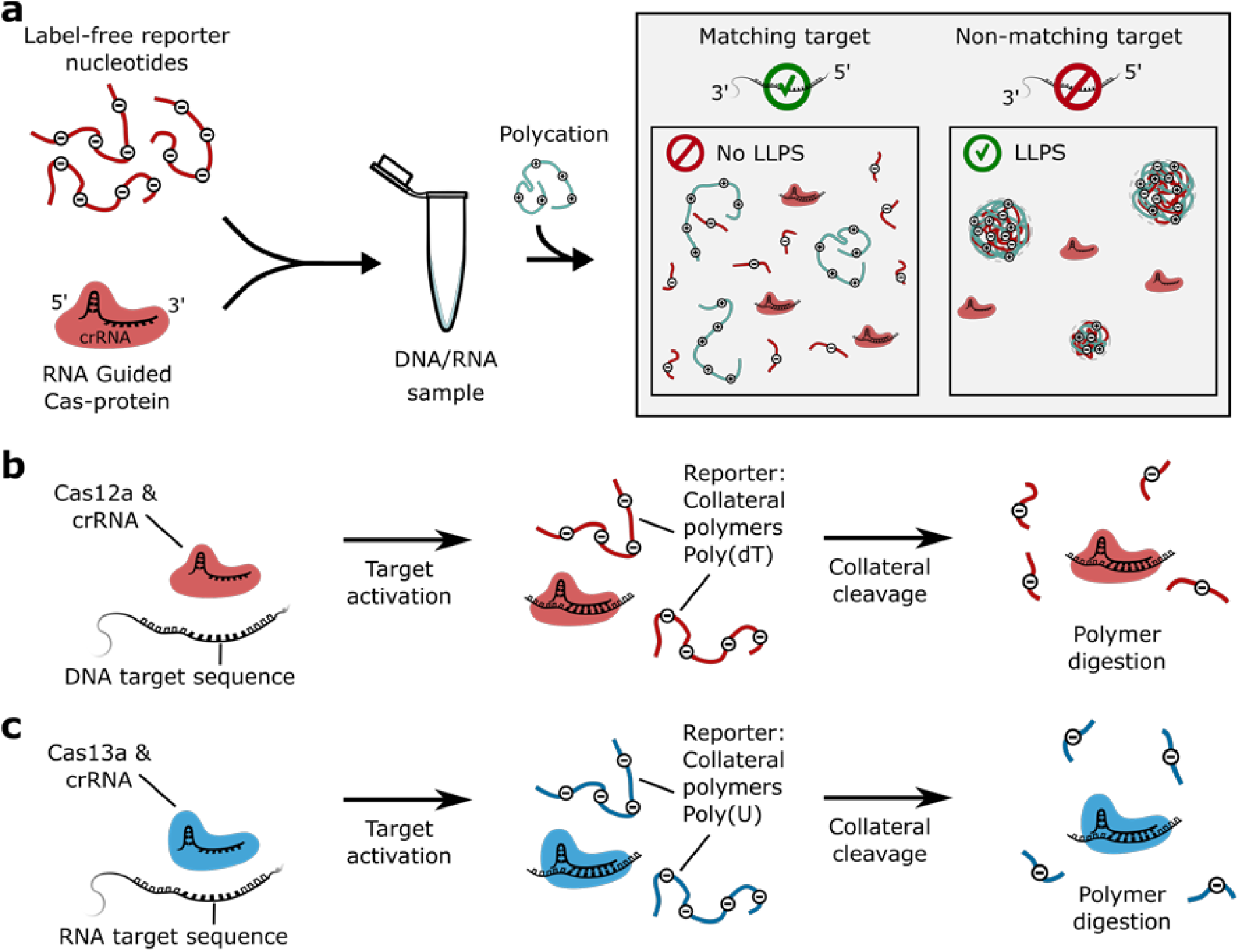
Molecular mechanisms of nucleotide detection via LLPS, Cas12a, and Cas13a. **(a)** Schematic for the detection of short DNA and RNA sequences via LLPS, which is facilitated by the collateral cleavage activity of CRISPR-Cas endonucleases towards single stranded DNA and RNA. RNA-guided endonuclease and collateral polymers are incubated with a sample. After incubation the mixture is complemented with a polycation to induce phase separation. In the presence of a matching target, the mixture will be transparent because collateral polymers are degraded, preventing LLPS. In the absence of target the mixture turns turbid due to LLPS of collateral polymer and the polycation. **(b)** RNA guided Cas12a targets DNA and is activated by a crRNA complementary single-stranded or double-stranded DNA target sequence. When active, the enzyme indiscriminately cleaves single stranded DNA, including poly(dT) oligonucleotides. **(c)** RNA guided Cas13a targets RNA and is activated by a crRNA complementary RNA target sequence, degrades RNA collaterally, with a preference for poly(U) (*11*).

To overcome the requirement of fluorescence-based detection, we have developed a detection method based on liquid-liquid phase separation (LLPS) (*15*–*17*). We exploit the fact that solutions of long nucleic acid polymers and positively charged polyelectrolytes can undergo liquid-liquid phase separation into a polymer-rich and a polymer-depleted phase. This process is also known as complex coacervation (*18, 19*). LLPS increases solution turbidity, such that it is visible to the naked eye (*20*). By using this effect in combination with the activity of CRISPR-Cas nucleases, it is possible to robustly couple the detection of specific DNA and RNA sequences to the turbidity of a sample solution. Our findings are explained by a mathematical model, which reveals a polymer length-dependent mechanism of liquid-liquid phase separation. We validate this theoretical result by liquid-liquid phase separation experiments with charged single stranded poly(dT) DNA.

## Results

### Inferring Cas12a and Cas13a activity from solution transparency/turbidity

To develop the molecular detection of nucleotide sequences, we tested the potential of LLPS as a readout mechanism for CRISPR-based nucleotide detection. A turbidity-based DNA and RNA nucleotide detection was implemented by the following in vitro experiments. We mixed crRNA-guided Cas12a, with and without complementary single stranded DNA target, and a 60-mer of poly(dT) (T60). When the positively charged polyelectrolyte poly-L-lysine (pLL, 15-30 kDa) was added immediately after mixing, both solutions turned turbid, indicating the presence of poly(dT) through visible liquid-liquid phase separation (**Fig. 2a**, left column). However, the solution containing the target remained transparent (**Fig. 2a**, right column) when pLL was added after 60 minutes incubation, whereas the solution without target still turned turbid. Thus, solution transparency indicates the successful detection of DNA targets.

**Figure 2:**
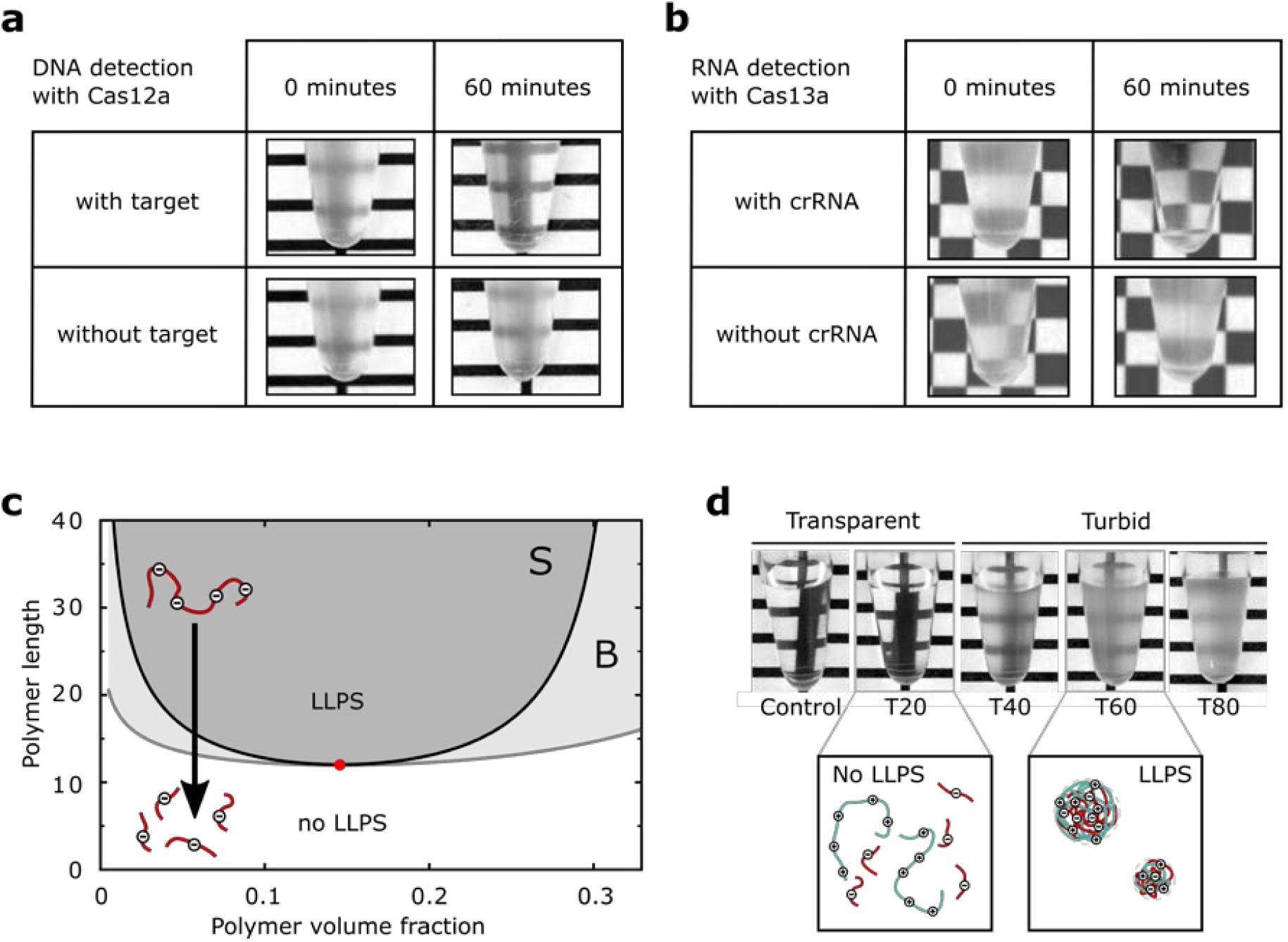
CRISPR-based sequence detection using LLPS. Cas12a and Cas13a activity can be assessed in a turbidity assay by exploiting phase separation of suitable oligonucleotides and positively charged polyelectrolytes. **(a)** Cas12a activity as observed from the absence of turbidity. The enzyme was incubated with poly(dT) (T60), both with and without a target sequence. Phase separation was induced with poly-L-Lysine. **(b)** Cas13a activity could be inferred by inducing LLPS of poly(U) and spermine. Cas13a was incubated with target, poly(U), and with or without crRNA. **(c)** Phase separation requires polymers longer than a minimum length for a given polymer volume fraction. The three regions in the diagram correspond to homogeneous solution (white, no LLPS), a metastable regime in region B (light grey, above the binodal curve), and the regime of spontaneous phase separation in region S (dark grey, above the spinodal curve). The red dot indicates the critical point of the system. If polymers are shorter than the critical length as indicated by the critical point, phase separation does not occur, independent of polymer volume fraction. Cleavage of polymers into pieces prevents phase separation (as indicated by the black arrow), which is why Cas12a and Cas13a can disrupt the phase separating behaviour of the solution. For simplicity the presented phase diagram is for symmetric polymer solutions based on Flory-Huggins (*24, 25*) theory and the Voorn-Overbeek (*19*) model (see Methods for details). **(d)** LLPS of poly(dT) and pLL depends on poly(dT) length. The pictures show transparent and turbid mixtures with different poly(dT) lengths ranging from T20 to T80. Long poly(dT) (>40 nt) leads to turbid and robustly phase separating solutions with 5 mg/mL pLL, whereas solutions with short poly(dT) (<40 nt) are transparent and do not phase separate (at equal volume fraction of nucleotides).

The above approach is also feasibility for sequence specific RNA detection. We identified a system composed of poly(U)-RNA and the tetravalent polycation spermine (*20*) as a suitable combination for liquid-liquid phase separation of RNA. We incubated Cas13a, the RNA target, and poly(U), with and without crRNA. When spermine was added immediately after mixing the components, both solutions turned turbid (**Fig. 2b**, left column). However, when spermine was added after 60 minutes, the sample that lacked crRNA underwent liquid-liquid phase separation and turned turbid, whereas the sample that contained crRNA remained clear (**Fig. 2b**, right column), indicating the successful activation of collateral RNase activity and prevention of LLPS. The result shows that LLPS-based detection is also compatible with RNA detection through Cas13a.

Having established that liquid-liquid phase separation of poly(dT) and poly(U) oligonucleotides with polycations can be exploited for CRISPR-based nucleic acid detection, we wanted to better understand the observed phase separation phenomenon and its underlying biophysical mechanism.

### Oligonucleotide length determines the onset of LLPS

Whether polymer solutions undergo liquid-liquid phase separation, i.e., spontaneously demix or not, depends both on the polymer concentration and polymer length (*21*–*23*). This means that there must be a critical polymer length above which demixing occurs at a given polymer concentration, whereas below a critical polymer length the solution components are uniformly mixed. In order to better understand the aforementioned nucleotide detection principle, we sought to identify the critical polymer length at which LLPS occurs.

To this end, we developed a thermodynamic model for solutions of oppositely charged polymers (see Methods) (*19, 24, 25*). The model predicts that in general, one can find conditions where a solution with long polymers phase-separates, whereas a solution with short polymers does not phase-separate, but will stay uniformly mixed. This prediction is shown by the phase diagram of polymer solutions as a function of polymer volume fraction and polymer length (**Fig. 2c**).

To test our theoretical prediction experimentally, we studied mixtures of various lengths of poly(dT) at constant volume fraction and the positively charged peptide poly-L-lysine. We found that phase separation robustly occurred for poly(dT) longer than 40 nucleotides and that phase separation was absent for shorter poly(dT). Notably, the change in turbidity of the mixtures could easily be distinguished by eye (**Fig. 2d**), which supports our observation that a shortening of poly(dT) through Cas12a endonuclease activity is preventing solution turbidity arising from LLPS.

In summary, the theory of charged polymer solutions can explain the phase behaviour of different poly(dT) oligonucleotides (**Fig. 2d**) and why Cas-proteins activity can prevent LLPS. Initially, the polymer length exceeds the critical length for phase separation. However, when degradation through Cas-proteins progresses and the average nucleic acid polymer length falls below the phase boundary (binodal curve in **Fig. 2c**), the solution loses its ability to phase-separate. Thus, the absence of detectable phase separation (*i.e.* solution clarity) indicates the Cas12a and Cas13a protein activity.

## Discussion

We have found a method for the label-free detection of short nucleotide sequences through transparency/turbidity of sample solutions by using CRISPR-Cas nucleases Cas12a and Cas13a. Instead of fluorescent reporter molecules, we chose oligonucleotides that undergo liquid-liquid phase separation to enable solution turbidity as a readout. We used crRNA-guided Cas12a or Cas13a to detect nucleotide target sequences and we exploited the enzymes’ collateral degradation activity towards nucleic acid polymers to affect the solution turbidity. For the readout, we complemented the solution with polycations to trigger LLPS of nucleic acid polymers. In case of successful target detection the collateral polymer has been digested, LLPS is not possible, and the solution remains transparent. If no target has been recognized the solution turns turbid, because interactions between nucleic acid polymers and the added polycations lead to LLPS and increased solution turbidity.

We have shown a proof-of-concept for the detection of DNA and RNA sequences through solution transparency/turbidity by combining programmable CRISPR-Cas nucleases Cas12a and Cas13a with liquid-liquid phase separation of oligonucleotides. Furthermore, we propose a mathematical model that is consistent with our results and explains how LLPS of interacting polymers depends on polymer length. Experiments where we used single-stranded poly(dT) primers of various sizes and poly-L-lysine confirmed the length-dependent transition between phase-separated and homogeneously mixed polymer solutions.

There is a variety of molecular mechanisms that allow for LLPS under specific conditions (*16*), and polymer length has been recognized as one of the important factors (*21*). Simple model systems, such as the one presented here, will help to obtain a better understanding of LLPS also in the cell biological context (*17*). As a recent example consider the cyclic GMP–AMP synthase, which forms liquid condensates with double stranded DNA depending on DNA length (*26*).

The above insights regarding polymer length-dependent LLPS helped us to devise the nucleotide detection assay based on CRISPR-Cas nucleases with limited equipment requirements. Existing nucleotide detection applications with Cas12a and Cas13a use chemically labelled nucleotides, fluorescence detection equipment, or the implementation on a paper strip. Because in our case transparency and turbidity of solution are the readout of the assay, it is possible to assess the test outcome with the unaided eye or a simple device to measures optical solution properties. We tested the limits of LLPS-based nucleotide detection with Cas12a and Cas13a. In our conditions the target detection limits were in the µM (ssDNA) and nM (RNA) range, which is comparable to reported values in the absence of amplification steps (*4*) for both enzymes (**Fig. S3**, Supporting Material). To enhance the detection limit for diagnostic applications for example, the target sample can be amplified using standard methods such as PCR, RPA, or T7 RNA polymerase, which increases the sensitivity by orders of magnitude (*3*–*5*).

Lastly, we explored the possibility to combine the nucleotide detection reaction and the triggering of LLPS. However, pLL as well as spermine interfered with the CRISPR-Cas proteins’ endonuclease activity and inhibited nucleotide detection, which leads us to speculate that metabolites like spermine may inhibit collateral cleavage in eukaryotic cells. This view is supported by the observation that Cas13a collateral cleavage does not occur *in vivo* in eukaryotic cells, since activated Cas13a neither compromised cell growth nor did it affect the RNA size distribution (*27*).

We anticipate that the presented LLPS-based nucleotide-detection assays will be useful in various high-throughput or low-cost point-of-care diagnostic applications, especially where the availability and operability of high-end laboratory equipment is limited.

## Methods

### Theoretical model

An important and well-known feature of LLPS in polymer solutions is that phase separation depends on the polymer length (*21, 22*). Liquid-liquid phase separation occurs when the energetic penalty for mixing the two liquids is larger than the entropic gain of that mixing.

In this section, we present a mathematical model for the free energy of the solution, from which we can calculate the phase diagram of the solution, and find under which conditions it will undergo phase separation.

Following the seminal work by Flory and Huggins (*24, 25*), a polymer solution can be described using a lattice model, where each lattice site is occupied by one of the components from the solution (*15, 16*). The size of a lattice site is chosen to be equal to the molecular volume of the solvent *v*. We denote the molecular volume of each component *i* by *V*_*i*_, and its valency by *Z*_*i*_. By using a mean-field approximation, one obtains an expression for the free energy in which only the volume fractions of polymers in solution are important. Based on the Flory-Huggins model (*24, 25*) and its extension to charged polymers by Voorn and Overbeek (*19*), the free energy per unit volume *f* (in units of *k*_*B*_*T*) of a polymer solution with *n* components (including the solvent) can be written as (*21*)

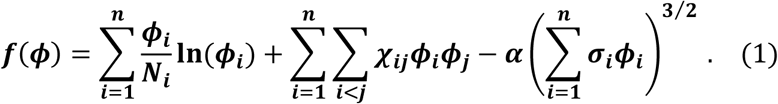

For component *i, ϕ*_*i*_ denotes the volume fraction, *N*_*i*_ = *V*_*i*_/ *v* denotes the effective chain length, *χ*_*ij*_ is the Flory-Huggins interaction parameter between components *i* and *j, α* is the electrostatic interaction parameter (*α ≈* 3.7 for aqueous solutions at room temperature (*28*)) and *σ*_*i*_ = *Z*_*i*_/*N*_*i*_ is the charge density. The first term of the free energy represents the entropy of mixing of the solution, and contains an explicit dependence on the effective chain length of each component. Due to its inversely proportional relation with polymer length, this term rapidly loses significance for long polymers. The second term represents the Flory-Huggins non-ionic interactions, and the third term represents the electrostatic interactions as in the Voorn-Overbeek model. For a qualitative demonstration of how polymer length influences phase separation behaviour, we make the simplifying assumption that the solution is symmetric and contains no added salts. The three components of the system are then the solvent molecules (index *s*) and two oppositely charged polymer species (indexes 1 and 2) of equal polymer length *N*_1_ = *N*_2_ = *N*; by definition *N*_*s*_ = 1. We further simplify our expressions by assuming that the two polymer species have equal charge density *σ*_1_ = *σ*_2_ = *σ*, equal polymer volume fraction 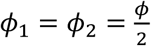, and that the only relevant Flory-Huggins interaction is that between the polymers and the solvent, so that *χ*_1*s*_ = *χ*_2*s*_ = *χ* and *χ*_12_ = 0. In this case, the free energy can be written as

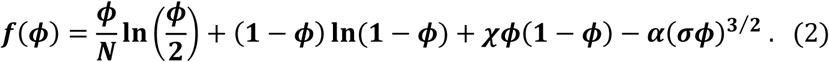

The free energy determines the thermodynamic properties of the polymer solution, where the first two terms represent the entropy of mixing, and the last two terms represent energetic contributions due to polymer-solvent and electrostatic interactions between the polymer species. The system will exhibit a uniform mixed phase if *f*(*ϕ*) is a convex function, and phase separate for any concave regions of *f*(*ϕ*). For fixed values of *χ* and *α, f*(*ϕ*) is convex below a critical value of *N*, but becomes concave for longer polymers. In **Fig. S4** (Supporting Material), the free energy and the chemical potential (μ = *∂f*/*∂ϕ*) are shown for effective chain lengths larger than, equal to, and lower than the critical length. We also calculated the phase diagram shown in **Fig. 2c** of the main text from the free energy given in equation (2) above. The critical point (*ϕ*_*c*_, *N*_*c*_), indicated by the red dot in **Fig. 2c**, can be computed from the condition that both the second and the third derivative of the free energy vanish

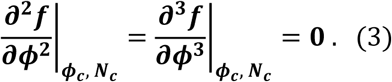

The critical point indicates the smallest polymer length *N*_*c*_ for which the solution can exhibit phase separation. The spinodal curve (black line in **Fig. 2c**) is given by the condition that the second derivative of the free energy vanishes

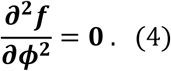

Inside the spinodal curve the solution is unstable and thus always phase separates, yielding stable coexistence of a polymer rich liquid phase and a polymer poor liquid phase. The volume fractions of the coexisting phases are indicated by the binodal curve (grey line in **Fig. 2c**), which is given by the condition that the chemical potential and osmotic pressure of each component are equal in both phases. To find the binodal, we (numerically) solve the simultaneous equations

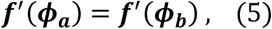

and

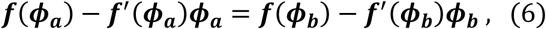

for *N* ≥ *N*_*c*_. Bet ween the bin odal and spi nodal cur ves, the coe xistence of two liq uid phases is metastable and the solution needs to overcome an energy barrier to demix. For the diagram in **Fig. 2c**, we set the parameters to *χ* = 0.5, *σ* = 0.22, *α* = 3.655. In the abs ence of the ele ctrostatic ter m, Flory-Huggins theory predicts LLPS to occur above a critical interaction strength (*16*) *χ*_*c*_ = 2 for *N* = 1. However, for *N* > 1, *χ*_*c*_ decr eases with incr easing *N*, and the polym er solut ion stays homogenous for polymer lengths of up to *N ≈* 10^4^ for *χ* = 0.5. The Voorn-Overbeek term in the free energy thus contributes the necessary interaction strength for phase separation to occur in the parameter regime we have chosen.

### Poly(dT) cleavage gel

To assess the collateral activity of Cas12a on T60 ssDNA with gel-electrophoresis (**Fig. S1a**), Cas12 was loaded with crRNA by mixing ∼8.6 µM Cas12a, 10 µM crRNA (Table S1) in 1x C12RB buffer and incubating for 15 minutes at 37 °C. Additionally, five tubes were prepared, each containing 1 µM T60 and 200 nM target DNA in 1x C12RB. Loaded Cas12a was added to each of the tubes and left for time intervals 1, 15, 30, 45 and 60 minutes. The reactions were stopped by flash-freezing the reaction tubes in liquid nitrogen, and addition of RNA gel loading dye followed by heating to 70 °C for 10 minutes and quickly cooling on ice for 5 minutes prior to running the samples on a non-denaturing poly acrylamide gel.

### Turbidity assays

Cas12a was loaded and activated as for the poly(dT) cleavage assay (see above). For the turbidity assay displayed in **Fig. 2a**, the four tubes displayed contained 187 nM of the activated Cas12a, 22 µM T60 in 1.1x C12RB buffer. The tubes shown in the upper row of **Fig. 2a** furthermore contained 220 nM DNA target. pLL was directly added to the tubes in the left column, to a final concentration of 0.5 mg/mL. The tubes in the right column were first incubated for one hour at 37 °C before pLL was added.

The turbidity assay for Cas13a (**Fig. 2b**) consisted of ∼50 nM Cas13a, ∼0.3 ng/µL Cas13a target and 0.1 wt% poly(U) mixed in 1.1x C13RB. The tubes in the upper row, also contained ∼0.3 ng/µL crRNA for Cas13a (Table S1). The tubes in the left column contained spermine from the start of the experiment at a final concentration of 1.0 wt%, whereas the tubes shown in the right column were first incubated at 37 °C for one hour before spermine was added.

### Buffer conditions and chemicals

Cas13a reactions as well as RNA/spermine experiments were carried out in C13RB buffer (40 mM Tris-HCl, 60 mM NaCl, 6 mM MgCl_2_, pH 7.3). Cas12a reactions and poly(dT)/pLL experiments were carried out in C12RB buffer (20 mM Tris-HCl, 66.6 mM KCl, 5 mM MgCl_2_, 1 mM DTT, 5% (v/v) glycerol, 50 µg/mL heparin, pH 7.5).

Poly(U) (polyuridylic acid potassium salt), spermine (spermine tetrahydrochloride), pLL (poly-L-lysine hydrobromide, 15-30 kDa), Trizma hydrochloride, sodium chloride and magnesium dichloride were bought from Sigma-Aldrich. Poly(dT) oligonucleotides T20, T40, T60, T80 were bought from Ella Biotech. RNase Alert, RNA Gel loading dye and Nuclease free water were bought from Thermo Fisher. DNase Alert was bought from Integrated DNA Technologies. RPA kit was bought from TwistDx.

### Fluorescence assays

We used fluorescence assays to test for collateral cleavage activity of Cas12a (**Fig. S1b**) with target (blue line), and without target sequence (orange line). Fluorescence measurements were done (*λ*_*ex*_ = 534 nm, *λ*_*em*_ = 585 nm) in a Tecan Infinite M200 PRO plate-reader. Cas12a was loaded with crRNA for a minimum of 15 minutes in 1x C12RB before the measurements, both reactions were done at 37 °C and as triplicate.

The fluorescence assay for Cas13a (**Fig. S2**) consisted of 50 nM Cas13a mixed with reaction buffer (1x C13RB), 0.3 ng/µL target RNA, 0.3 ng/µL crRNA, and 125 nM RNase Alert. Fluorescence (*λ*_*ee*_ = 490 nm, *λ*_*ee*_ = 520 nm) was recorded every three minutes using a Synergy H1 (BioTek) fluorimeter. All measurements were done at 37 °C and as triplicate.

### Protein expression and purification

#### Cas12a purification

The plasmid pET21a encoding Cas12a (*7*) (AsCpf1) was transformed into Rosetta *E. coli* cells, which were grown on agar plate with 100 µg/mL ampicillin and 25 µg/mL chloramphenicol. Cells were grown in LB with addition of 100 µg/mL ampicillin and 25 µg/mL chloramphenicol, to an OD_600_ of 0.6 (37°C, 180 rpm). Cultures were put on ice for 30 minutes, and Cas12a production was induced by the addition of 1 mM IPTG. Cultures were left for 16 hours at 18 °C with rotation (180 rpm). Subsequent steps were done at 4°C. Cultures were spun down at 7000 g for 10 minutes, and the supernatant was discarded. Pellets were resuspended in lysis buffer (50 mM NaH_2_PO_4_, 500 mM NaCl, 1 mM DTT, 10 mM imidazole, pH 8.0) with addition of 1 protease inhibitor tablet per 50 mL of lysis buffer. Cells were disrupted in a French Pressure Cell at 1000 bar twice and spun down at 16,000 g for 30 min. Supernatant was filtered with a 0.45 µm pore size, applied to His-Select Affinity resin and incubated for 30 minutes. Resin was spun down at 2000 g for 1 minute, and the supernatant was discarded. The resin was loaded onto a gravity column and washed twice with wash buffer (50 mM NaH_2_PO_4_, 500 mM NaCl, 1 mM DTT, 30 mM imidazole, pH 8.0). Cas12a was eluted with elution buffer (50 mM NaH_2_PO_4_, 500 mM NaCl, 1 mM DTT, 250 mM imidazole, pH 8.0). The recovered protein was spun down at 14,000 g for 10 min and the supernatant was loaded onto a HiLoad 16/600 Superdex 200 PG column. Fractions of 1 mL were recovered and the protein content measured at OD280. Protein containing samples were pooled and the elution buffer exchanged with exchange buffer (20 mM Hepes, 150 mM KCl, 10 mM MgCl_2_, 1% glycerol, 0.5 mM DTT, pH 7.5) using a Millipore Amicon 10,000 NMWL centrifugal filter unit. Sample was spun down at 14,000 g for 10 min, and the supernatant snap frozen in fractions for further use. The final protein concentration was determined using a Bradford assay.

#### Cas13a purification

The plasmid (*3*) pC013 encoding Cas13a was transformed into BL21(DE3) *E. coli* cells, which were grown on agar plate with 100 µg/mL ampicillin. Cells were grown in Terrific Broth (TB) medium, containing 12 g/L tryptone, 24 g/L yeast extract, 9.4 g/L K_2_HPO_4_ and 2.2 g/L KH_2_PO_4_, with addition of 100 µg/mL ampicillin, to an OD_600_ of 0.6 (37 °C, 180 rpm). Cultures were put on ice for 30 minutes, and subsequently Cas13a production was induced by the addition of 0.5 mM IPTG. Cultures were left for 16 hours at 18 °C with rotation (180 rpm). Subsequent steps were done at 4 °C. Cultures were spun down at 5200 g for 15 minutes, and the supernatant was discarded. Pellets were resuspended in 1xPBS buffer, and centrifuged (3220 g, 10 minutes). Pellets were resuspended in lysis buffer (20 mM Tris-HCl, 0.5 M NaCl, 1 mM DTT, 25 mM imidazole, pH 8.0) with addition of 100 u/mL benzonase, 0.25 mg/mL lysozyme and 1 protease inhibitor tablet per mL. Cells were French pressed (3 rounds at 100 kbar) and cell lysate was pelleted for 45 minutes at 16,000 g. Supernatant was filtered with a 0.45 µm pore size, applied to His-Select Nickel Affinity gel and incubated for one hour with rotation. Resin was spun down (1 minute, 2000 g), and the supernatant was discarded. The resin was loaded onto a gravity column, and washed with lysis buffer. Cas13a was eluted with elution buffer (20 mM Tris-HCl, 0.5 M NaCl, 1 mM DTT and 250 mM imidazole, pH 8.0), and dialyzed overnight against storage buffer (50 mM Tris-HCl, 0.6 mM NaCl, 5% v/v glycerol, 2 mM DTT, pH 7.5). Final protein concentration was determined using a Bradford assay.

### Nucleic acid preparations

#### Targets for Cas12a and Cas13a

ssDNA target for Cas12a assays was ordered from Ella Biotech (5’-Cga gta aca gac atg gac cat cag ATC TAC AAC AGT AGA AAT TCT ATA GTG AGT CGT ATT ACT T-3’).

Target RNA for Cas13a assays was produced by Recombinant Polymerase Amplification (RPA, TwistDx) and simultaneous *in vitro* transcription of part of the tetracycline resistance gene from the plasmid pSB1T3. The target amplification was done by mixing 1 ng of plasmid with 1 µM forward primer (5’-AAT TCT AAT ACG ACT CAC TAT AGG gat gcc ctt gag agc ctt caa c-3’) containing a T7 promoter overhang, 1 µM reverse primer (5’-cct cgc cga aaa tga ccc a-3’), 4 µL Murine RNase inhibitor (NEB), 1 µL T7 polymerase (NEB), 5 mM MgCl_2_, 1 mM ATP, 1 mM CTP, 1 mM GTP and 1 mM UTP and RPA rehydration buffer. This was transferred into an RPA mix and 14 mM MgAc was added to start the reaction, which was left for three hours at 37 °C. Target RNA was purified using RNeasy MinElute (Qiagen), quantified using NanoDrop, and stored at −80 °C.

#### crRNA preparation

crRNA for Cas12a was bought from IDT. crRNA for Cas13a, guided towards RNA target, was prepared by annealing of template DNA oligonucleotides and *in vitro* transcription. Primers 5’-AAT TCT AAT ACG ACT CAC TAT AGG GGA TTT AGA CTA CCC CAA AAA CGA AGG GGA CTA AAA C-3’ and 5’-gcc gca ctt atg act gtc ttc ttt atc aGT TTT AGT CCC CTT CGT TTT TGG GGT AGT CTA AAT CCC CTA TAG TGA GTC GTA TTA GAA TT-3’ were mixed in 1 µM concentration and annealed by heating to 95 °C for 2 minutes and gradually cooling to 50 °C in steps of 5 °C every ten seconds, and then cooling to 30 °C in ten seconds. The resulting annealed DNA was *in vitro* transcribed in a mix containing 1 mM ATP, 1 mM CTP, 1 mM GTP, 1 mM UTP, 1 µL T7 polymerase, 2 µL murine RNase inhibitor and RNA polymerase buffer, to a final volume of 100 µL. Reaction was left at 37 °C for 3 hours, and cleaned using RNeasy MinElute. The concentration was determined using NanoDrop.

## Acknowledgements

We thank members of the iGEM team 2017 from Delft University of Technology for their contributions: Aafke van Aalst, Kimberley Barentsen, Kelly Hamers, Floor de Jong, Fiona Murphy, Guillermo Serena Ruiz, Gabriella Tany, Isabell Trinh, Jasper Veerman, Amária Vledder and Hielke Walinga, and team supervisors Aljoscha Wahl, Cristóbal Almendros Romero, Anthony Birnie, Hirad Daneshpour, David Foschepoth, Patrick de Jonge, Sebastian Kieper, Benjamin Lehner, Rebecca McKenzie, Jochem Vink and Esengül Yildirim. The plasmids encoding Cas12a and Cas13a were a kind gift from the Feng Zhang lab (Broad Institute). This work was supported by The Netherlands Organization for Scientific Research (NWO/OCW), as part of the Frontiers of Nanoscience program, and the Fonds Ernest Solvay managed by the Koning Boudewijnstichting, Franklin L. Nobrega was supported by a NWO Veni grant 016.Veni.181.092, Stan J.J. Brouns was supported by the European Research Council (ERC Starting Grant 639707) and a TU Delft start-up grant, and L. Reese was supported by FOM programme nr. 110 (NWO).

## Author contributions

L.R., W.K.S., and S.J.J.B. conceived the study; W.K.S., J.M.J., L.R., F.L.N., and A.C.H. carried out the experiments; W.K.S. and T.I., developed the theoretical model with input from L.R.; W.K.S., F.L.N., M.D., T.I., S.J.J.B., and L.R. analysed and discussed the data; W.K.S. and L.R. wrote the paper with input from all authors.

## Materials & Correspondence

Correspondence should be addressed to Stan Brouns.

