## Supporting Material for "CRISPR-based DNA and RNA detection with liquid-liquid phase separation"

**Cas12a/Cas13a cleavage activity.** We assessed the non-specific cleavage activity of activated Cas12a on a poly(dT) 60-mer substrate. Degradation of poly(dT) was observed immediately (~1 min) after the addition of activated Cas12a (**Fig. S1a**), suggesting that poly(dT) is an excellent substrate to Cas12a. In a complementary fluorescence-based assay we observed cleavage activity of activated Cas12a on the timescale of minutes (**Fig. S1b**). The assay measured nuclease activity by fluorescence arising from a DNA-linked fluorophore-quencher pair. Upon cleavage of the DNA link between fluorophore (F) and quencher (Q) the fluorophore becomes fluorescent.

**Sensitivity of the method.** To assess the sensitivity of the detection method, we tested the target detection limit of the method for both DNA and RNA detection (**Fig. S3**). We obtained a visible readout at sub-micromolar concentrations of ssDNA target using Cas12a, and at picomolar concentrations of RNA target using Cas13a. Both values are similar to previous reports in the literature in the absence of amplification steps[4]. By including appropriate amplification reactions attomolar sensitivities are possible when T7 RNA polymerase[3] or recombinase polymerase amplification[5] are used with double stranded target DNA. We expect that our method can be adapted for such high detection sensitivities when needed by using T7 or RPA amplification, because phase separation depends on polymer length in a binary manner.

**Figure S1**

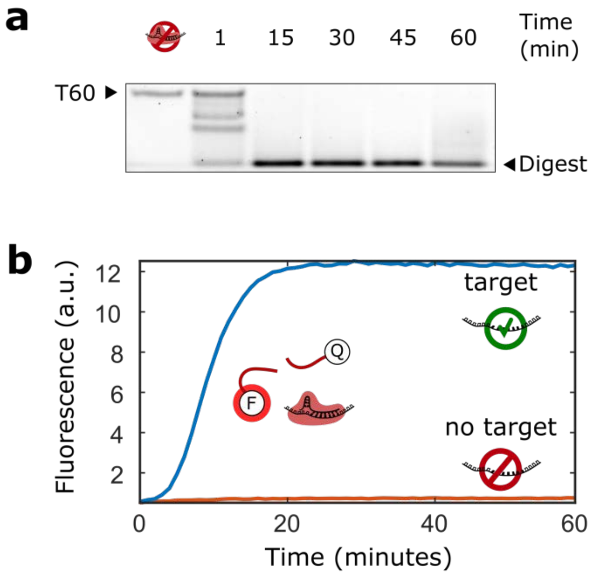


**Figure S1: Cas12a cleavage activity.**

**(a)** Cas12a cleaves poly(dT). The gel shows degradation of a poly(dT) 60-mer (T60) in the presence of activated Cas12a over time.

**(b)** The activity of Cas12a in the presence of ssDNA target in a fluorescence (DNase Alert, IDT, see Methods for details). The curves show DNase Alert fluorescence with (blue) and without (orange) target.

**Figure S2**

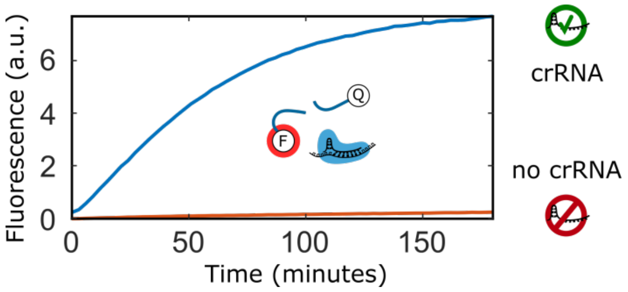


**Figure S2: Cas13a cleavage activity.**

Fluorescence-based assay confirming that in the presence of target, Cas13a activity depends on the presence of crRNA (see Methods). The curves represent fluorescence intensity from RNase Alert of solutions with (blue) and without (orange) crRNA.

**Figure S3**

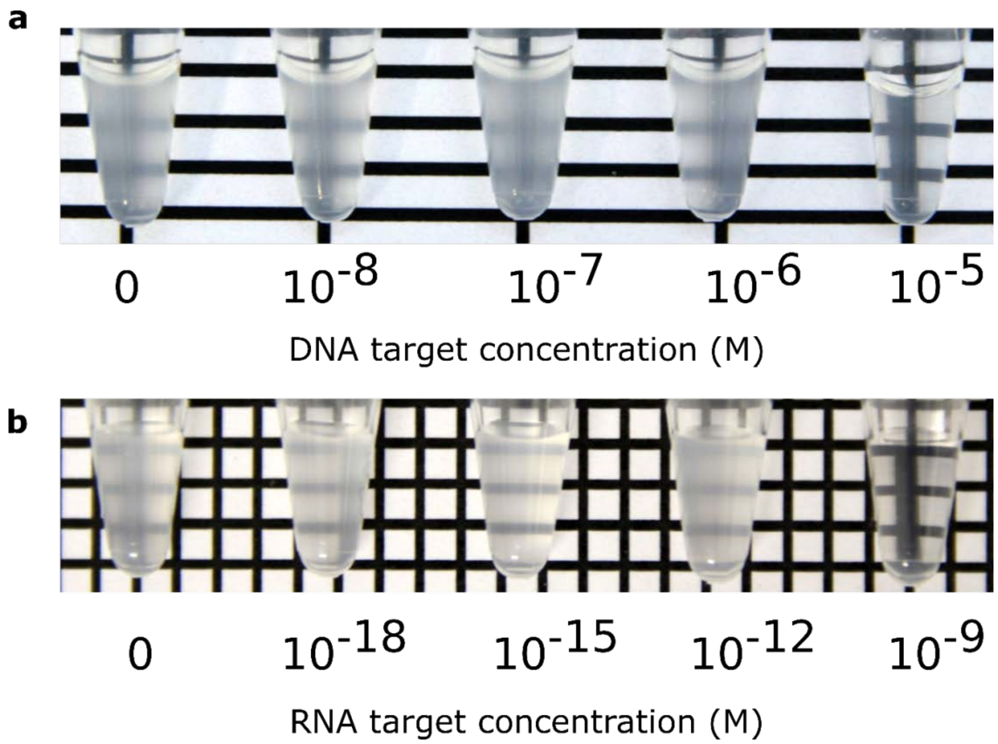


**Figure S3: Detection limit of the phase-separation assay for Cas12a and Cas13a are in the micromolar and nanomolar range, respectively.**

**(a)** All tubes initially contained ~189 nM activated Cas12a, 22 µM T60 in 1.1x C12RB buffer, and varying target concentrations (molar concentrations indicated below the tubes). After 1 hour of incubation at 37°C, pLL was added to a final concentration of 0.5 mg/mL.

**(b)** All tubes contained ~37 nM Cas13a, 0.11 wt% poly(U), 45 nM crRNA, C13RB buffer and varying concentrations of RNA target (molar concentrations indicated below tubes). After 3 hours of incubation at 37 °C, spermine was added to a final concentration of 1.0 wt%.

**Figure S4**

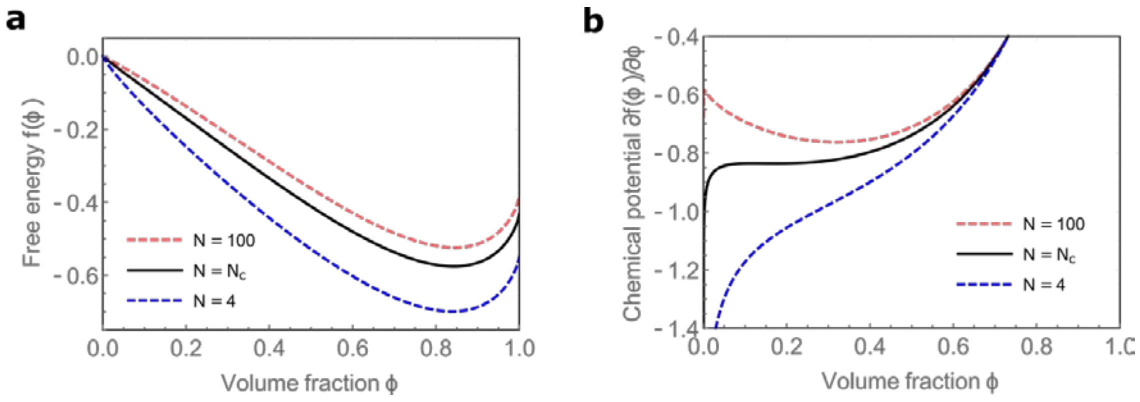


**Figure S4: The dependence of the free energy of a symmetric solution of charged polymers for different effective chain lengths.**

**(a)** Free energy as a function of volume fraction $\phi$ for effective polymer chain lengths $N$ lower than, equal to and larger than the critical length ($N_c \approx 12$). The parameters for this plot are the same as for the phase diagram in **Fig. 1c** ($\chi = 0.5$, $\sigma = 0.22$, $\alpha = 3.655$).

**(b)** The chemical potential is the derivative of the free energy as displayed in panel (a). For effective polymer chain lengths larger than the critical length, a local minimum appears in the chemical potential. For effective polymer chain lengths lower than the critical length, there is no local minimum.

**Table S1: Sequences of crRNA used for Cas12a and Cas13a.**

| crRNA | Sequence (5'->3') with target complementarity (lower case) |
|---|---|
| Cas12a | UAAUUUCUACUCUUGUAGAUcugaugguccaugucuguuacuc |
| Cas13a | GGGGAUUUAGACUACCCCAAAAACGAAGGGGACUAAAACugauaaag aagacagucauaagugcggc |
